# Adsorption of respiratory syncytial virus (RSV), rhinovirus, SARS-CoV-2, and F+ bacteriophage MS2 RNA onto wastewater solids from raw wastewater

**DOI:** 10.1101/2023.05.04.539429

**Authors:** Laura Roldan-Hernandez, Alexandria B. Boehm

## Abstract

Despite the wide adoption of wastewater surveillance, more research is needed to understand the fate and transport of viral genetic markers in wastewater. This information is essential for the interpretation of wastewater surveillance data and the development of mechanistic models that link wastewater measurements to the number of individuals shedding virus. In this study, we examined the solid-liquid partitioning behavior of four viruses in wastewater: SARS-CoV-2, respiratory syncytial virus (RSV), rhinovirus (RV), and F+ coliphage/MS2. We used two approaches to achieve this: we (1) conducted laboratory partitioning experiments using lab-grown viruses and (2) examined the distribution of endogenous viruses in wastewater. Partition experiments were conducted at 4°C and 22°C; wastewater samples were spiked with varying concentrations of each virus and stored for three hours to allow the system to equilibrate. Solids and liquids were separated via centrifugation and viral RNA concentrations were quantified using reverse-transcription-digital droplet PCR (RT-ddPCR). For the distribution experiment, wastewater samples were collected from six wastewater treatment plants and processed without spiking exogenous viruses; viral RNA concentrations were measured in wastewater solids and liquid. Overall, RNA concentrations were higher in solids than the liquid fraction of wastewater by approximately 3–4 orders of magnitude. Partition coefficients (K_F_) from laboratory experiments were determined using the Freundlich model and ranged from 2,000–270,000 ml·g^-1^ across viruses and temperature conditions. Distribution coefficients (K_d_) determined from endogenous wastewater viruses were consistent with results from laboratory experiments.Further research is needed to understand how virus and wastewater characteristics might influence the partition of viral genetic markers in wastewater.

**Synopsis:** We examined the solid-liquid partitioning behavior of SARS-CoV-2, RSV, RV, and F+coliphage/MS2 RNA in wastewater influent. Overall, partition/distribution coefficients were similar across viruses and temperature conditions.

## Introduction

Multiple countries are currently monitoring the spread of COVID-19 by measuring the genetic markers of SARS-CoV-2 variants in wastewater and primary settled solids (hereafter referred to as wastewater matrices). A few wastewater surveillance programs also monitor the genetic markers of common respiratory diseases like influenza, respiratory syncytial virus (RSV), rhinovirus (RV), and human metapneumovirus (HMPV).^1–5^ This information can be used by public health officials to monitor infection trends, complement clinical surveillance data, and strengthen public health responses.^6^ Despite the widespread adoption of wastewater surveillance, research is still needed to understand the fate and transport of viral genetic markers in wastewater matrices. This information is essential for linking wastewater surveillance data to the number of individuals in sewersheds shedding virus, and disease incidence and prevalence. ^7–9^

Viral adsorption can be influenced by the physical, chemical, and biological characteristics of wastewater (e.g., temperature, pH, organic matter) and characteristics of viruses (e.g., virus structure and size).^10–13^ Previous studies suggest that viruses and their genetic markers tend to partition more favorably to the solid fraction of wastewater matrices than the liquid fraction.^2,14–17^ For example, Mercier et al.^2^ studied the distribution of endogenous influenza A virus (IAV) in wastewater influent and primary sludge and found that the majority of IAV RNA was in settled solids compared to suspended solids (larger than 0.45 µm) and the liquid fraction of these matrices. Li et al.^16^ also examined the distribution of endogenous SARS-CoV-2 RNA (N1, N2, and E gene targets) in wastewater influent and found that the majority of viral genetic markers were in the solid fraction of wastewater. A few studies have also reported higher concentrations of viral genetic markers in primary sludge compared to paired wastewater influent samples. For instance, Wolfe et al.^18^ found that IAV RNA concentrations were 1,000 times higher in primary settled solids than wastewater influent on mass equivalent basis. Similar results have been reported for SARS-CoV-2, MPOX virus, and pepper mild mottle virus (PMMoV) RNA where viral RNA concentrations were enriched by 3-4 orders of magnitude in primary settled solids compared to wastewater influent.^14–16,19^ Yin et al.^20^ summarized the solid-liquid distribution of different strains of enteroviruses, hepatitis A, adenovirus, rotavirus, and bacteriophages in wastewater and activated sludge and found that viral adsorption can vary greatly between viruses and wastewater matrices. Still, in all cases, viruses tended to partition to wastewater solids.

A few studies have also examined the equilibrium and kinetic adsorption of viruses and their genetic markers in wastewater matrices. For example, Ye et al.^12^ studied the sorption kinetics of four infectious lab-grown human virus surrogates (MHV, ϕ6, MS2, and T3) in wastewater influent and found that enveloped viruses partitioned more to the solid fraction of wastewater compared to non-enveloped viruses. A similar study was conducted by Yang et al.^11^ but using molecular methods (quantitative PCR) to quantify the concentrations of four lab-grown surrogates (Phi6, MS2, T4, and Phix174) in activated sludge. Partition coefficients (converted from log K_F_; also known as the Freundlich coefficient) were 4.1×10^6^, 5.4×10^5^, 1.2×10^5^, and 8.5 ×10^3^ mL·g^-1^ for Phi6, MS2, T4, and Phix174 in sludge, respectively. Yin et al.^20^ also measured the sorption of human Adenovirus 40 (HAV40) in primary and secondary sludge and found that the majority of HAV40 DNA was adsorbed into the solids fraction of these matrices. Partition coefficients (reported as K_p_ in the paper) were 3.7×10^4^ mL·g^-1^ and 4.0×10^4^ mL·g^-1^ in primary and secondary sludge, respectively. Researchers have also examined the equilibrium and kinetic adsorption of SARS-CoV-2 RNA onto passive samplers designed for wastewater surveillance.^21^

In this study, we examined the partitioning behavior of four viruses in wastewater: SARS-CoV-2, respiratory syncytial virus (RSV), rhinovirus (RV), and MS2/F+ coliphage. We achieve this through laboratory partitioning experiments and through examination of the distribution of these viruses in actual wastewater samples. SARS-CoV-2, RSV, and RV were chosen for the study because their equilibrium partitioning and distribution in wastewater has not been previously studied. MS2 was chosen because it is widely used as a surrogate for pathogenic respiratory viruses in lab experiments. These viruses represent both enveloped (SARS-CoV-2 and RSV) and non-enveloped (RV and MS2) viruses. Additionally, the human pathogenic viruses chosen in this study are targets for wastewater-based epidemiology monitoring efforts. Understanding the partitioning behavior of viral genetic markers could inform wastewater sampling strategies and help optimize methods for processing wastewater and primary sludge samples. Partition and distribution coefficients can also help inform complex mathematical models that aim to estimate or predict the number of positive cases in communities.^22^

## Materials and Methods

### Overview

We conducted two sets of experiments to examine the partition and distribution, respectively, of SARS-CoV-2, RSV, RV, and F+ coliphage in wastewater influent. The characteristics of these viruses are shown in Table 1. The partitioning experiment was conducted using lab-grown SARS-CoV-2, RSV-A, RV-B, and MS2. For these experiments, wastewater influent samples were spiked with varying concentrations of each virus and incubated at two different temperatures (4°C and 22°C) to allow the system to equilibrate. After incubation, influent samples were centrifuged and decanted to obtain an aliquot from the liquid and solid fractions. RNA was extracted from the aliquots and quantified using reverse-transcription-digital droplet PCR (RT-ddPCR). The distribution experiment examined the distribution of endogenous SARS-CoV-2, RSV, RV, and F+ coliphage in actual wastewater samples. Influent samples were collected from six wastewater plants and processed using the previously described method, but without spiking with viral surrogates. The following sections provide a detailed description of the experiments. Reporting of methods follows EMMI guidelines^23^ (Figure S1 provides EMMI checklist and details).

**Table 1:** Characteristics of SARS-CoV-2, RSV, RV, and MS2^35^.

| **Virus** | **Family/Genus** | **Genome Type** | **Structure** | **Shape** | **Genome Size (kb)** | **Virion Size (nm)** |
| --- | --- | --- | --- | --- | --- | --- |
| SARS-CoV-2 | Coronaviridae | + ssRNA | enveloped | spherical | 30 | 50 –140 |
| RSV | Pneumoviridae | - ssRNA | enveloped | spherical | 15 | 150 –250 |
| Rhinovirus | Picornavirus | + ssRNA | nonenveloped | icosahedral | 7 | 15 – 30 |
| MS2 | Leviviridae | + ssRNA | nonenveloped | icosahedral | 3.6 | 23 – 28 |

**Table 2:**
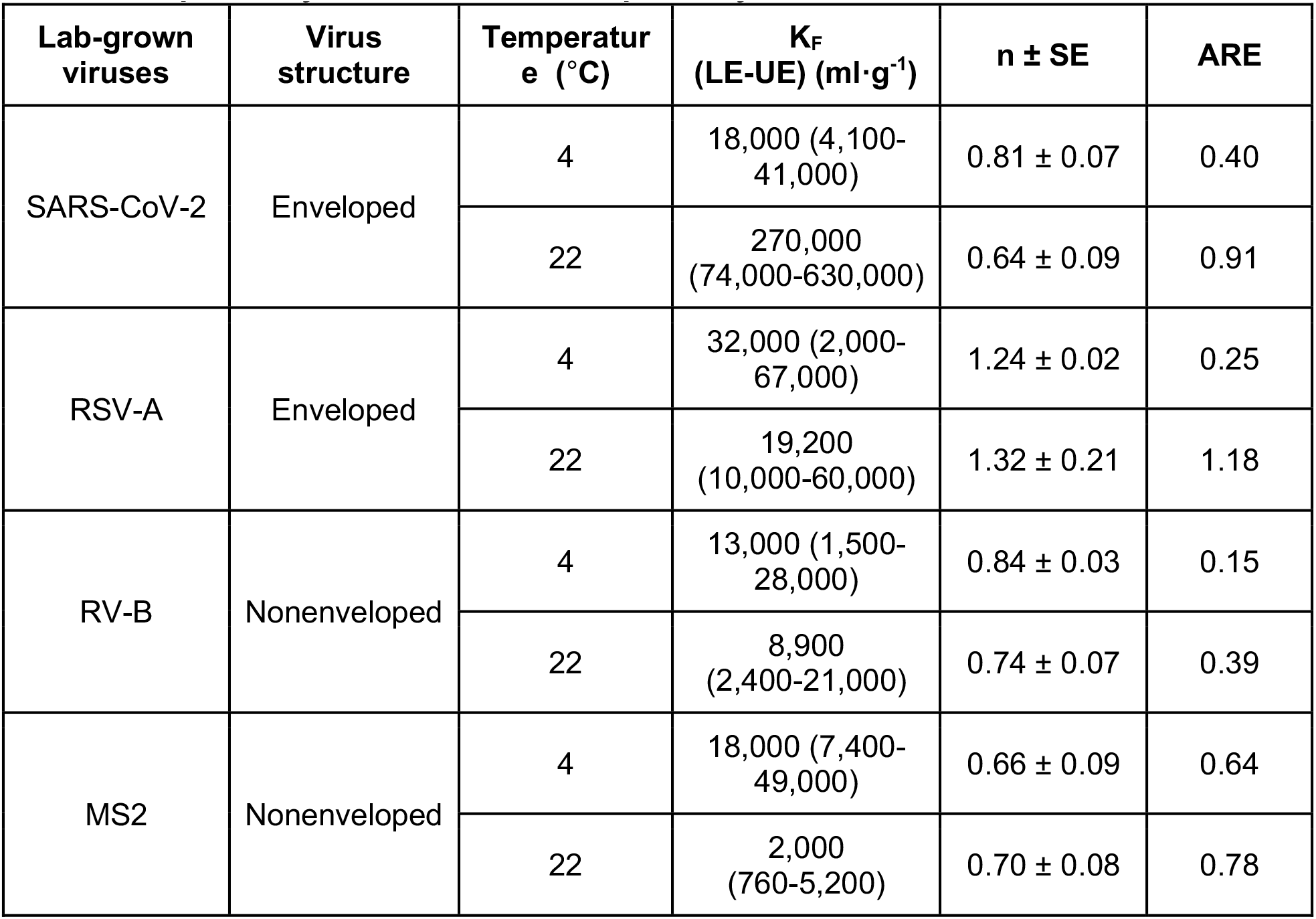
Isotherm parameters (K_F_ and n) and average relative error (ARE) of Freundlich models for the adsorption of SARS-CoV-2, RSV-A, RV-B, and MS2 in wastewater at 4°C and 22°C; SE, LE, and UE are the standard error, the lower SE bound, and the upper SE bound as reported by lm function in R, respectively. n and ARE are dimensionless.

### Wastewater sample collection

The partitioning experiment was conducted using two influent samples from the Palo Alto Regional Water Quality Control Plant (PA). The plant serves approximately 215,000 people and treats an annual average daily flow of 19.8 million gallons per day (MGD). Total suspended solids (TSS) and pH levels range from 220–360 mg/L and 7.5–7.8, respectively. We collected approximately two liters of a 24-hour influent composite sample on September 20, 2022 for the 4°C experiment and on October 28, 2022 for the 22°C experiment. Samples were collected in 10% HCl acid-washed plastic containers, stored at the respective experimental temperatures, and spiked with a mixture of lab-grown SAR-CoV-2, RSV-A, RV-B, and MS2 within 24 hours of sample collection (see detailed methods below) for experiments.

### Virus purification and spike cocktails

Heat-inactivated SARS-CoV-2 (Isolate: USA-WA1/2020; catalog no. 0810587CFHI), RSV-A (catalog no. 0810040ACF), and RV-B (catalog no. 0810284CF) were purchased from ZeptoMetrix (Buffalo, New York). The manufacturer inactivates SARS-CoV-2 by heating the virus at 60°C for 1 hour. RSV-A and RV-B are viable viruses suspended in cell culture fluids. *Escherichia coli* phage MS2 (DMS no. 13767) was purchased from the DSMZ German Collection of Microorganisms and Cell Cultures. Viruses were purified to remove viral culture fluid using Amicon® Ultra-0.5 ml centrifugal filters (100 kDa MWCO; Millipore UFC5100) following the manufacturer’s instructions. Briefly, 0.5 ml of virus stock, as received from the vendor, was added to individual centrifugal filters and centrifuged at 14,000xg for 5 min. The filters were immediately flipped and centrifuged at 1,000xg for 2 minutes to recover the retentate. A series of dilutions were prepared using autoclaved phosphate-buffered saline (PBS; Fisher BioReagents, Pittsburgh, Pennsylvania) to achieve a viral genome concentration of approximately 1×10^3^, 1×10^4^, 1×10^5^, 1×10^6^, and 1×10^7^ cp/µl PBS. A total of five spike cocktails were prepared by mixing equal volumes of purified heat-inactivated SARS-CoV-2, RSV-A, RV-B, and MS2 stock. The final concentrations of the five stock cocktails ranged from approximately 1×10^3^–1×10^7^ cp/µl PBS (10^3^, 10^4^, 10^5^, 10^6^, 10^7^) for each virus.

### Preanalytical processing

Wastewater samples were thoroughly mixed by inverting 3–4 times and aliquoted into eighteen 50 ml centrifuge tubes (hereafter, referred to as subsamples). Five sets of three subsamples were then spiked with one of the five different concentrations of spiked cocktails to achieve a final concentration of approximately 10^3^, 10^4^, 10^5^, 10^6^, or 10^7^ cp/ml for each virus. Three additional subsamples were reserved to measure the background concentration of endogenous SARS-CoV-2, RSV, RV, and F+ coliphage RNA. After spiking, subsamples were stored at 4°C or 22°C, depending on the experiment, and gently mixed (∼20 rpm) using a tube roller (Globe Scientific, GSCI-GTR-AVS) for approximately three hours to allow the system to equilibrate. The time needed to reach equilibrium (< 3 hours) was determined based on a preliminary experiment (see Figure S2 and S3) and it is consistent with other virus adsorption studies conducted in wastewater.^11,12^ After three hours, subsamples were centrifuged at 24,500xg for 20 minutes at the temperatures of the experiments. This process removes solid particles with hydrodynamic radii greater than 0.3 µm in diameter.^24^ 200 µl of the supernatant was transferred to a 2 ml collection tube and spiked with 5 µl of bovine coronavirus vaccine (BCoV; Zoetis, CALF-GUARD®; Parsippany-Troy Hills, NJ) as an extraction recovery control. The BCoV vaccine comes lyophilized and was resuspended in 3 mL of molecular-grade water. These aliquots represent the liquid fraction of the wastewater sample.

The remaining supernatant was decanted and the pellet represents dewatered solids. A portion of the dewatered solids was reserved to calculate the dry weight. Dewatered solids were weighed before and after drying at 105°C for 24 hours. To collect solids for viral analysis, approximately 0.1 g of dewatered solids were collected from the bottom of the centrifuge tube and aliquoted into 2 ml microcentrifuge tubes using a disposable spatula (Fisher Scientific, catalog no. 50-476-569). Dewatered solids were resuspended in BCoV-spiked DNA/RNA shield (Zymo Research, catalog no. R1100-250) at a concentration of approximately 75 mg/ml; this concentration of solids in the buffer has been shown to alleviate downstream RT-ddPCR inhibition.^25^ Spiked DNA/RNA shield was pre-prepared using 1.5 μl of BCoV per ml of DNA/RNA shield. Three to five grinding balls (OPS DIAGNOSTICS, GBSS 156-5000-01) were added to a 2 ml microcentrifuge tube and the mixture was homogenized at 4 m/s for 1 minute using the MP Bio Fastprep-24TM (MP Biomedicals, Santa Ana, CA). The homogenized aliquots were then centrifuged for 5 min at 5,250 x g and 200 µl of the supernatant was transferred to a 2 ml collection tube.This aliquot contains viral targets from the solid fraction of the wastewater sample. Liquid and solid aliquots were stored at 4°C overnight and the nucleic acids were extracted from them the next day.

### RNA extraction

RNA was extracted from the solid and liquid aliquots using the Qiagen AllPrep PowerViral DNA/RNA kit and further purified using the Zymo OneStep PCR inhibitor removal columns (Zymo Research, Irvine, CA) following the manufacturer’s instructions. RNA extracts were aliquoted into 1.5 ml DNA LoBind tubes, stored at -80°C for less than two weeks, and thawed once (1 freeze-thaw cycle) before quantification. Nuclease-free water and BCoV-spiked DNA/RNA shield were used as negative and positive extraction controls, respectively. These controls were carried through the extraction process with one set of positive and negative controls per extraction batch (∼15 solid or liquid aliquots/batch of extraction).

### RNA Quantification

SARS-CoV-2 and RSV were quantified using a duplex assay described in Hughes et al.^26^ and RV, MS2, and BCoV were quantified using singleplex assays from previous studies.^1,15,27^ Primers and probes were purchased from Integrated DNA Technologies (IDT, San Diego, CA) and are provided in Table S1. Using NCBI Blast, we determined that the MS2 assay will detect, in addition to MS2, the following sequenced and deposited strains of genotype group 1 (GI) F+ RNA coliphages: JP501, M12, DL16, DL52, and DL54 which could potentially be present in wastewater. There could be additional unsequenced/undeposited endogenous wastewater F+ RNA coliphages that are also detectable using this assay. Therefore, detections with the MS2 in wastewater will be referred to as F+ coliphage, hereafter.

All viral targets were quantified using the One-Step RT-ddPCR Advanced Kit for Probes (Bio-Rad 1863021). For the RT-ddPCR, 20 µl of a 22 µl reaction mix was prepared for each well. The reaction consisted of 5.5 µl of template RNA, 5.5 µl of Supermix, 2.2 µl of 200 U/µl Reverse Transcriptase (RT), 1.1 µl of 300 mM dithiothreitol (DDT), 4.4 µl of nuclease-free water, and 3.3 µl of primer and probe mixture with a final concentration of 900 nM and 250 nM, respectively.

RNA extracts were processed undiluted and in duplicate (two technical replicates). Nuclease-free water and viral RNA extracts for each target were used as negative and positive PCR controls, respectively, and each run in three wells per plate. Positive controls were extracted from the SARS-CoV-2, RSV, RV, and MS2 purified stocks; which are the same virus stocks used to prepare the spike cocktails. Droplets were generated using the AutoDG Automated Droplet Generator (Bio-Rad, Hercules, CA) and amplified using the C1000 Touch™ Thermal Cycler (Bio-Rad, Hercules, CA). Thermal cycling conditions for each RT-ddPCR assay are shown in Table S2. After amplification, droplets were analyzed using the QX200 droplet reader and the Quantasoft Analysis Pro Software. Wells with less than 10,000 droplets were excluded and technical PCR replicates (wells) were merged before performing the dimensional analysis. Merged wells needed to have at least three positive droplets to be considered positive The estimated lower measurement limit for solids and liquid aliquots were 3,500 cp/g and 0.7 cp/ml, respectively.

### Distribution of endogenous viruses in wastewater

The second experiment examined the distribution of endogenous SAR-CoV-2, RSV, RV, and F+ coliphage between liquid and solid fractions in wastewater samples. Influent samples (∼1 L of a 24-hour composite sample) were collected from six wastewater treatment plants on November 18, 2022. Samples were stored at 4°C and processed within 24 hours. These plants are part of an ongoing wastewater surveillance program and include the following: Oceanside Water Pollution Control Plant (OS), Southeast Water Pollution Control Plant (SE), Silicon Valley Clean Water Wastewater Treatment Plant (SV), Sunnyvale Water Pollution Control Plant (SU), San Jose-Santa Clara Regional Wastewater Facility (SJ), and South County Regional Wastewater Treatment Plant (GI). The location, estimated population served, annual daily average flow, pH, and TSS for each plant can be found in Table S3. Figure S4 provides a map of the sewershed (area served) for each wastewater treatment plant.

Wastewater samples were thoroughly mixed, poured into 50 ml centrifuge tubes, and processed. Three subsamples were prepared for each wastewater treatment plant, for a total of 18 subsamples. Solid and liquid aliquots were obtained from the wastewater samples and nucleic acids were extracted. Viral targets were quantified from the aliquots using the methods described above for the partitioning experiments.

### Dimensional analysis, adsorption models, and statistical analysis

SARS-CoV-2, RSV, RV, and MS2/F+ coliphage RNA concentrations were expressed in units of copies per gram dry weight solids (cp/g) or per ml liquid (cp/ml) for the solid and liquid fractions of wastewater, respectively, using dimensional analysis. BCoV recovery, used as an extraction and inhibition control, was calculated for the solid and liquid fractions as follows:

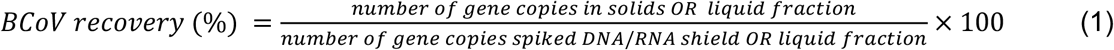

For the partition experiment using lab-grown viruses, 36 RNA concentrations (solids: N=18 and liquids: N=18) were obtained for each virus and isotherm experiment (4°C and 22°C). These concentrations represent triplicate measurements for six different initial conditions: one unspiked and five different spiked concentrations of viruses. For each virus and temperature condition, average viral RNA concentrations (solids: N=6 and liquids: N=6) and standard deviations were calculated across triplicate subsamples. To calculate the recovery of spiked viruses, we first subtracted the background concentrations of viral genetic markers from liquid and solid fractions of spiked subsamples and estimated the total RNA recovery in spiked subsample as follows:

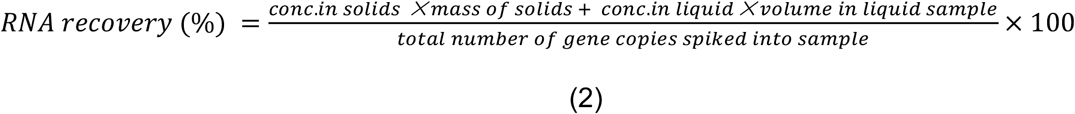

Partition experiment viral RNA concentrations in the solid and liquid fractions of wastewater were fit to Linear, Langmuir, and Freundlich isotherm models. These models have been used in previous virus partitioning studies^11,14,20,21^ and describe a multi-layer (Freundlich) or monolayer (Langmuir) adsorption process. A linear model is generally used when the coverage ratio of adsorption sites is minimal.^28^ The Linear, Freundlich, and Langmuir isotherm parameters were determined for each virus and temperature condition. The average relative error (ARE) was calculated for each model and compared to identify the model with the best fit. The Freundlich model produced the smallest ARE overall and therefore is discussed in the main paper (see Table S4 for results of other models). The nonlinear and linear forms of the Freundlich model are described as follows:

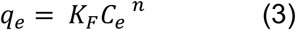

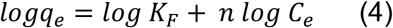

where q_e_ is the equilibrium concentration of viral genomes in solids (cp/g), C_e_ is the equilibrium concentration of viral genomes in the liquid fraction of wastewater, K_F_ is the Freundlich constant, and n is the adsorption intensity. A higher n indicates a stronger interaction between the absorbent (i.e., wastewater solids) and the adsorbate (i.e., spiked viruses). We determined the Freundlich isotherm parameters, logK_F_ (y-intercept) and n (slope), using a linear regression model (lm function) in R. Standard errors (SE) were also obtained from the linear regression model in R (summary function). Finally, we compared K_F_ obtained for the viruses under different temperature conditions by examining whether their values and standard deviations overlapped.

For the distribution experiment (examining endogenous viruses in wastewater samples), six RNA concentrations (solids: N=3 and liquids: N=3) were obtained for each virus and wastewater treatment plant. Only a few liquid fractions resulted in non-detects (ND). NDs were substituted with half of the lower measurement limit (0.35 cp/ml for concentrations measured in liquid fractions).The average concentration and standard deviation for each virus were calculated using data from the three replicate subsamples. The distribution coefficient was calculated as the ratio of average viral genome concentrations detected in the solid and liquid fractions of wastewater influent; K_d_=C_s_/C_w_; errors were determined by propagating errors on the numerator and denominator.We tested the null hypothesis that K_d_ were the same for endogenous viruses using a Kruskal-Wallis test; *p*<0.05 was used to assess statistical significance. Statistical analysis was performed in R (version 4.1.2). Finally, we compared the partition coefficients (K_F_) for spiked viruses at 4°C and 22°C to the distribution coefficients (K_d_) for endogenous viruses in wastewater for each virus to determine if K_F_ and K_d_ values and errors overlapped. We also summarized the results from previous experiments measuring the concentration of viral genetic markers in wastewater matrices to see if our results were consistent with previously reported K_F_ and K_d_ values (see Table 4).

**Table 4:**
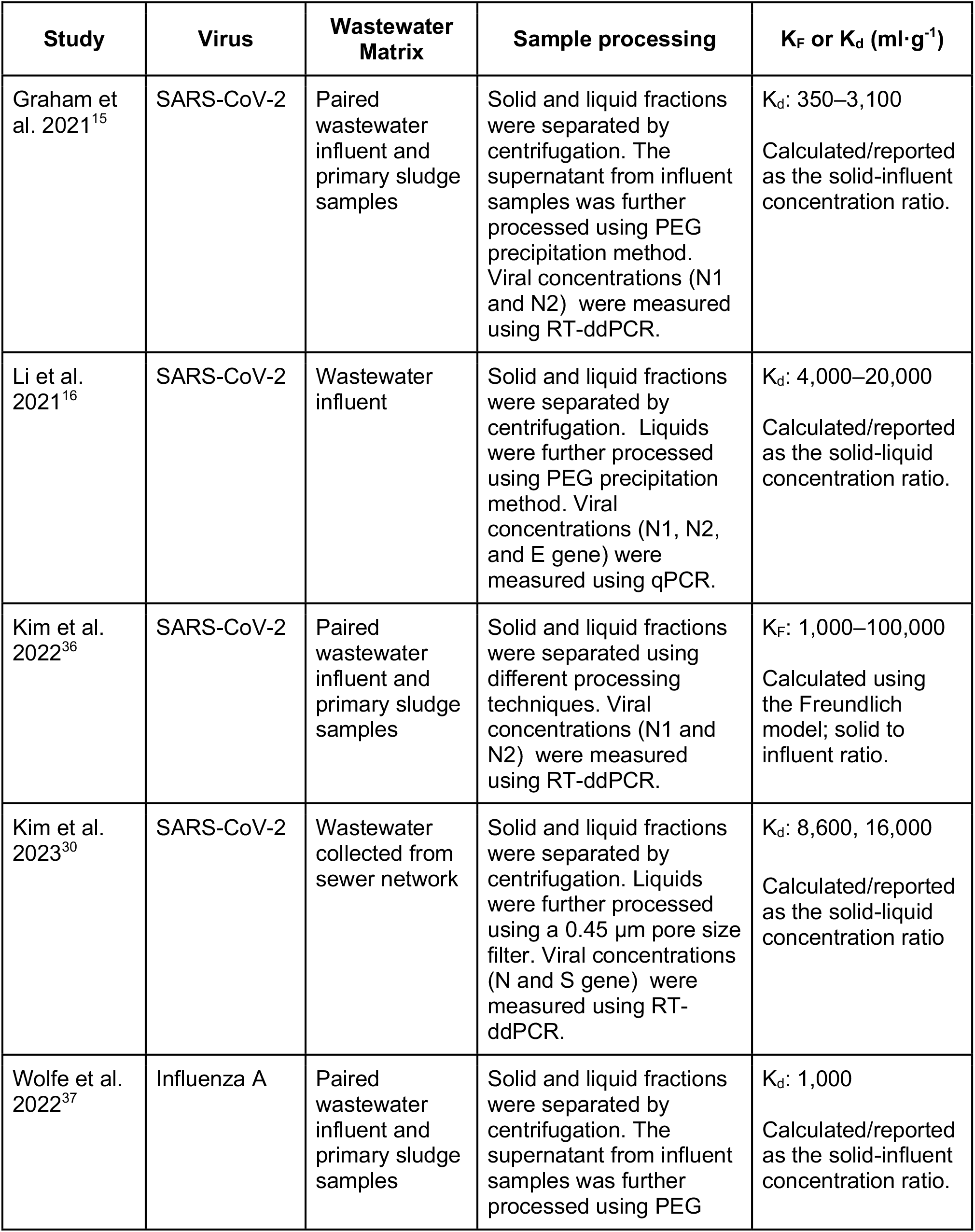
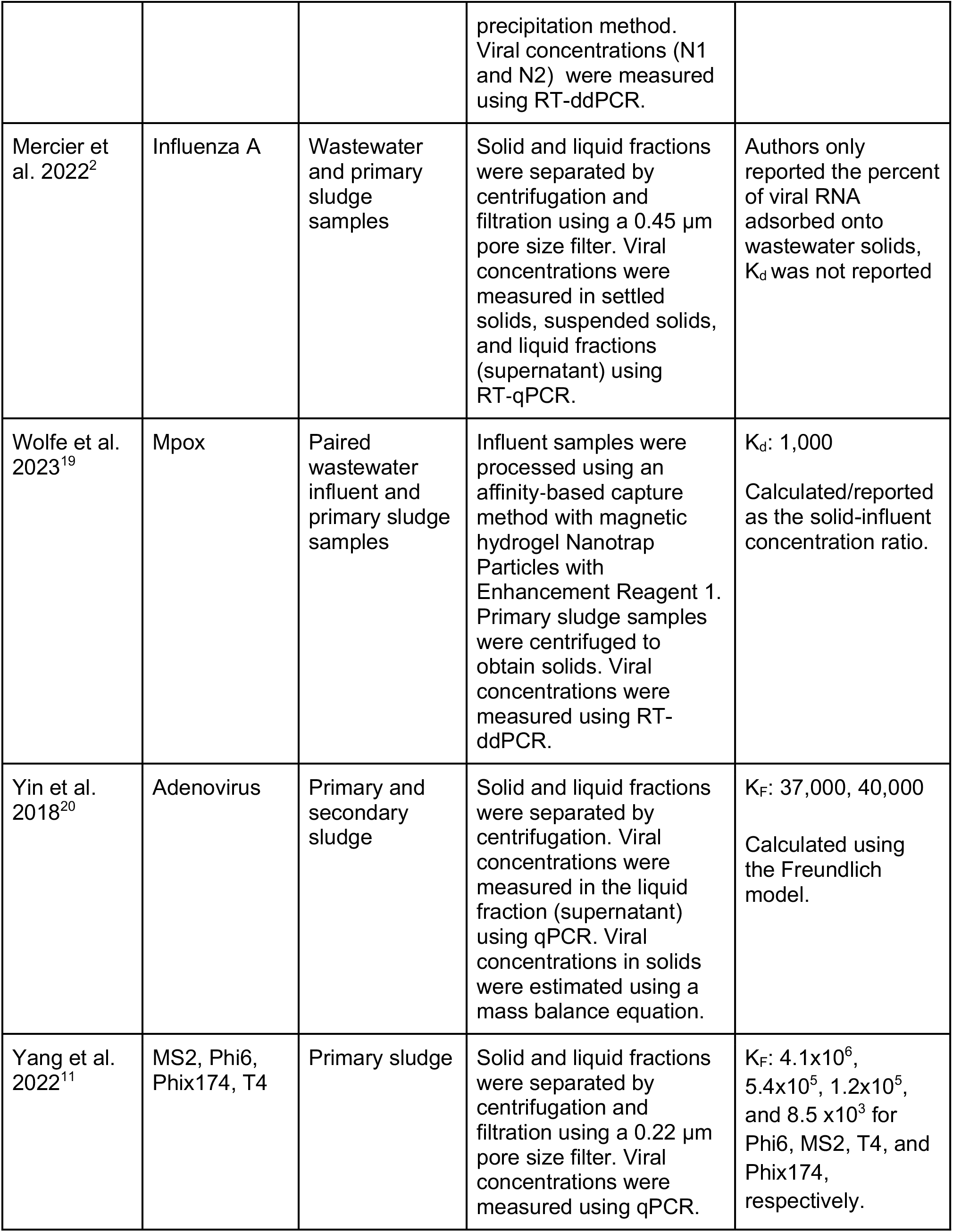
Results from previous experiments measuring the concentration of viral genetic markers in the solid and liquid fractions of wastewater matrices.

| Study | Virus | Wastewater Matrix | Sample processing | K<sub>F</sub> or K<sub>d</sub> (ml·g<sup>-1</sup>) |
| --- | --- | --- | --- | --- |
| Graham et al. 2021<sup>15</sup> | SARS-CoV-2 | Paired wastewater influent and primary sludge samples | Solid and liquid fractions were separated by centrifugation. The supernatant from influent samples was further processed using PEG precipitation method. Viral concentrations (N1 and N2) were measured using RT-ddPCR. | K<sub>d</sub>: 350–3,100<br>Calculated/reported as the solid-influent concentration ratio. |
| Li et al. 2021<sup>16</sup> | SARS-CoV-2 | Wastewater influent | Solid and liquid fractions were separated by centrifugation. Liquids were further processed using PEG precipitation method. Viral concentrations (N1, N2, and E gene) were measured using qPCR. | K<sub>d</sub>: 4,000–20,000<br>Calculated/reported as the solid-liquid concentration ratio. |
| Kim et al. 2022<sup>36</sup> | SARS-CoV-2 | Paired wastewater influent and primary sludge samples | Solid and liquid fractions were separated using different processing techniques. Viral concentrations (N1 and N2) were measured using RT-ddPCR. | K<sub>F</sub>: 1,000–100,000<br>Calculated using the Freundlich model; solid to influent ratio. |
| Kim et al. 2023<sup>30</sup> | SARS-CoV-2 | Wastewater collected from sewer network | Solid and liquid fractions were separated by centrifugation. Liquids were further processed using a 0.45 µm pore size filter. Viral concentrations (N and S gene) were measured using RT-ddPCR. | K<sub>d</sub>: 8,600, 16,000<br>Calculated/reported as the solid-liquid concentration ratio |
| Wolfe et al. 2022<sup>37</sup> | Influenza A | Paired wastewater influent and primary sludge samples | Solid and liquid fractions were separated by centrifugation. The supernatant from influent samples was further processed using PEG precipitation method. Viral concentrations (N1 and N2) were measured using RT-ddPCR. | K<sub>d</sub>: 1,000<br>Calculated/reported as the solid-influent concentration ratio. |
| Mercier et al. 2022<sup>2</sup> | Influenza A | Wastewater and primary sludge samples | Solid and liquid fractions were separated by centrifugation and filtration using a 0.45 µm pore size filter. Viral concentrations were measured in settled solids, suspended solids, and liquid fractions (supernatant) using RT-qPCR. | Authors only reported the percent of viral RNA adsorbed onto wastewater solids, $K_d$ was not reported |
| Wolfe et al. 2023<sup>19</sup> | Mpox | Paired wastewater influent and primary sludge samples | Influent samples were processed using an affinity-based capture method with magnetic hydrogel Nanotrap Particles with Enhancement Reagent 1. Primary sludge samples were centrifuged to obtain solids. Viral concentrations were measured using RT-ddPCR. | $K_d$: 1,000<br>Calculated/reported as the solid-influent concentration ratio. |
| Yin et al. 2018<sup>20</sup> | Adenovirus | Primary and secondary sludge | Solid and liquid fractions were separated by centrifugation. Viral concentrations were measured in the liquid fraction (supernatant) using qPCR. Viral concentrations in solids were estimated using a mass balance equation. | $K_F$: 37,000, 40,000<br>Calculated using the Freundlich model. |
| Yang et al. 2022<sup>11</sup> | MS2, Phi6, Phix174, T4 | Primary sludge | Solid and liquid fractions were separated by centrifugation and filtration using a 0.22 µm pore size filter. Viral concentrations were measured using qPCR. | $K_F$: $4.1 \times 10^6$, $5.4 \times 10^5$, $1.2 \times 10^5$, and $8.5 \times 10^3$ for Phi6, MS2, T4, and Phix174, respectively. |

## Results

### Extraction and PCR controls

Positive and negative extraction and PCR controls were positive and negative, respectively. BCoV was used as a process recovery and gross inhibition control. BCoV recoveries were similar between the solid and liquid fractions of wastewater. Solids and liquid aliquots had a median BCoV recovery of 0.40 and 0.35, respectively. Viral genome concentrations were not adjusted by recovery given the complexities associated with estimating recovery using surrogate viruses^29^ and given that the recoveries were similar. The total recovery of spiked SARS-CoV-2, RSV-A, RV-B, and MS2 RNA was approximately 35%, 45%, 22%, and 33%, respectively. These recoveries are similar to those of BCoV. Although inhibition can vary between assays, our previous work has determined there is minimal inhibition with the pre-analytical and analytical workflow,^25^ and the similar and high detection of BCoV and spiked viruses indicates inhibition is minimal.

### Partitioning of lab-grown viruses in wastewater influent

Endogenous SARS-CoV-2, RSV, RV, and MS2/F+ coliphage RNA were detected in the liquid and solid fractions of wastewater influent samples from PA (Table S5). Concentrations of endogenous virus make up 22%–95% of viral RNA in subsamples spiked with the lowest concentration of virus cocktail, but otherwise represent a negligible percentage of the RNA in subsamples spiked with a higher concentration of virus cocktail.

At equilibrium, the results from the partitioning experiments showed that RNA concentrations of spiked viruses were higher in solids than the liquid fraction of wastewater influent on a mass equivalent basis, by approximately 3–4 orders of magnitude. In the 4°C and 22°C isotherm experiments, q_e_ ranged from 1.3×10^4^–5.6×10^7^ cp/g for SARS-CoV-2, 4.2×10^3^–1.9×10^8^ cp/g for RSV, 2.3×10^4^–1.3×10^7^ cp/g for RV, and 4.1×10^3^–3.2×10^7^ cp/g for MS2. In the liquid fraction, C_e_ concentrations ranged from 0.1–1.3×10^3^ cp/ml for SARS-CoV-2, 0.2–4.5×10^2^ cp/ml for RSV, 1.6–4.8×10^4^ cp/ml for RV, and 0.8 –4.3×10^4^ cp/ml for MS2. The reported ranges represent measurements across spiking conditions and the two experimental temperatures.

Viral RNA concentrations (q_e_ and C_e_, see Figure 1) were fit to a Linear, Freundlich, and Langmuir model.The Freundlich isotherm models produced the lowest ARE compared to the other models based on the calculated partition coefficient (see Table S4 for results of Linear and Langmuir models). Table 3 shows the Freundlich isotherm parameters (K_F_ and n) for each virus and temperature condition. In the 4°C experiment, K_F_ and n ranged from 1.8×10^3^–3.2×10^3^ ml·g^-1^ and 0.66–1.24, respectively. Similar results were obtained in the 22°C experiments except for the partition coefficient of SARS-CoV-2. In the 22°C experiment, K_F_ and n ranged from 2.0×10^3^– 2.7×10^5^ ml·g^-1^ and 0.64–1.32, respectively. The partition coefficient of SARS-CoV-2 in the 22°C experiment was significantly higher (approximately one order of magnitude) compared to other viruses and temperature conditions. However, partition coefficients were not different across other viruses and temperatures (see Figure S5).

**Table 3:** Distribution coefficient (K_d_ =C_s_/C_w_) of endogenous SARS-CoV-2, RSV, RV, and F+ coliphage RNA in wastewater influent; SD is the standard deviation across triplicate subsamples.

| Wastewater treatment plant | $K_d \pm SD$ (ml·g<sup>-1</sup>) | | | |
| --- | --- | --- | --- | --- |
|  | SARS-CoV-2 | RSV | RV | F+ coliphage |
| Gilroy | 3,500 ± 820 | 1,300 ± 650 | 7,200 ± 1,400 | 490 ± 380 |
| San Jose | 11,000 ± 6,200 | 2,000 ± 760 | 16,000 ± 4,100 | 2,800 ± 1800 |
| Sunnyvale | 12,000 ± 7,600 | 1,800 ± 460 | 8,000 ± 4,800 | 950 ± 710 |
| SVCW | 5,600 ± 4,300 | 1,300 ± 450 | 11,000 ± 7,200 | 7,400 ± 6,300 |
| SEP | 4,400 ± 1,400 | 500 ± 320 | 2,100 ± 1,400 | 1,200 ± 760 |
| OSP | 3,000 ± 2,500 | 700 ± 470 | 2,800 ± 1,000 | 1,300 ± 470 |

**Figure 1:**
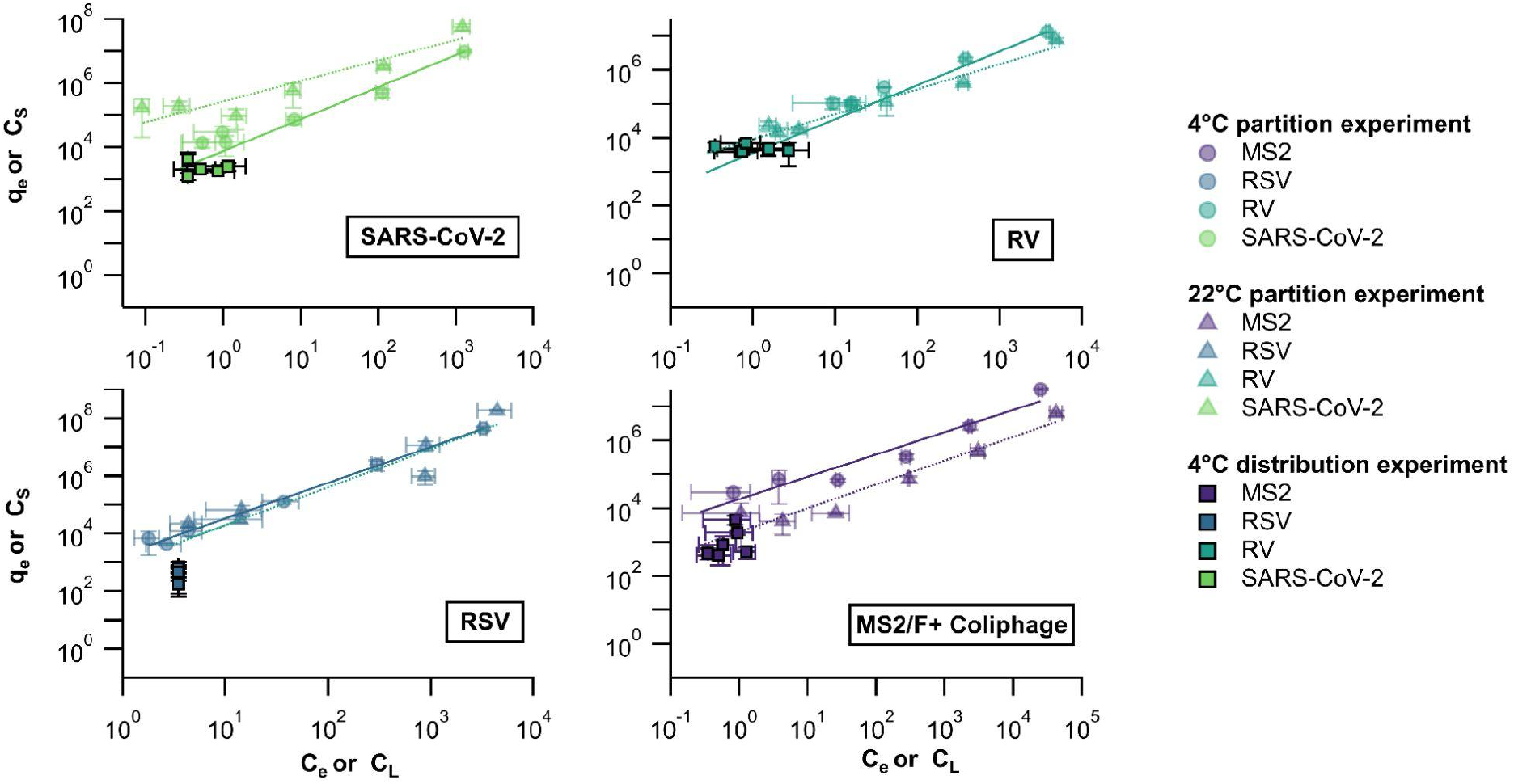
q_e_ and C_e_ from partition experiments at 4°C (circles) and 22°C (triangles) and C_S_ and C_L_ from distribution experiment at 4°C (squares, with black edges). Lines represent the Freundlich isotherm model and error bars represent the standard deviation across triplicate subsamples.

### Distribution of endogenous viruses in wastewater influent

In the wastewater of six wastewater treatment plants, we observed results similar to those obtained in the laboratory partitioning experiment. Viral RNA concentrations were higher in solids than the liquid fraction of wastewater, by approximately 3–4 orders of magnitude (see Figure 1). Across six wastewater treatment plants, C_s_ ranged from 1.6×10^3^–1.5×10^3^ cp/g (median = 3.2×10^3^ cp/g) for SARS-CoV-2, 9.1×10^2^–3.7×10^3^ cp/g (median = 1.7×10^3^ cp/g) for RSV, 2.1×10^3^–1.4×10^4^ cp/g (median = 6.0×10^3^ cp/g) for RV, and 4.9×10^2^–6.1×10^3^ cp/g (median = 1.1×10^3^ cp/g) for F+ coliphage. C_L_ ranged from 0.4–1.1 cp/ml (median = 0.4 cp/ml) for SARS-CoV-2, 0.4–2.7 cp/ml (median = 0.8 cp/ml) for RV, and 0.4–1.3 cp/ml (median = 0.7 cp/ml) for F+ coliphage. For RSV, C_L_ were ND across wastewater treatment plants; NDs were replaced with half of the lower measurable limit (0.35 cp/ml for viral concentrations in liquid fractions) to calculate the distribution coefficient (K_d_).

K_d_ was calculated as the ratio of C_s_/C_L_. Across wastewater treatment plants, K_d_ ranged from 3.5×10^3^–1.2×10^4^ ml·g^-1^ (median = 5.0×10^3^ ml·g^-1^) for SARS-CoV-2, 5.0×10^2^–2.0×10^3^ ml·g^-1^ (median = 1.3×10^3^ ml·g^-1^) for RSV, 2.1×10^3^–1.6×10^4^ ml·g^-1^ (median = 7.6×10^3^ ml·g^-1^) for RV, and 4.9×10^2^–7.4×10^3^ ml·g^-1^ (median = 1.2×10^3^ ml·g^-1^) for F+ coliphage (see Table 4). Overall, RV had the largest solid-liquid distribution, followed by SARS-CoV-2, F+ coliphage, and RSV. Note that K_d_ for RSV could be higher, but we were only able to estimate a lower bound for Kd since the measurement in liquid was ND. RV K_d_ was statistically higher from RSV and F+ coliphage K_d_ (Kruskal-Wallis and Dunn’s post hoc test, both *p*<0.05).

## Discussion

This is the first batch experiment examining the solid-liquid partitioning of SARS-CoV-2, RSV, and RV in wastewater and the first experiment to examine the distribution of endogenous RSV and RV in this matrix. Overall, higher concentrations of viral RNA were observed in solids compared to the liquid fraction of wastewater, for all viruses and temperature conditions; viral RNA concentrations were higher in solids by 3–4 orders of magnitude on a mass equivalent basis. Partition and distribution coefficients were also similar across viruses and temperature conditions, with K_F_ and K_d_ ranging from 490 ml·g^-1^–270,000 ml·g^-1^. Our results are consistent with previously reported partition/distribution coefficients for viral genetic markers in wastewater. For example, Li et al.^16^ measured the distribution of endogenous SARS-CoV-2 RNA (N1, N2, and E genes) in wastewater samples and found higher concentrations of SARS-CoV-2 RNA in solids compared to the liquid fraction; K_d_ (reported in their paper as the solid-liquid concentration ratio) ranged from 4,000–20,000 ml·g^-1^. Kim et al.^30^ also studied the distribution of endogenous SARS-CoV-2 RNA (S and N genes) in wastewater samples from two K-12 schools and observed similar results; viral RNA concentrations were higher in solids by three orders of magnitude and K_d_ (reported in their paper as the concentration ratio in solid to liquid samples) were 8,600 ml·g^-1^ and 16,000 ml·g^-1^ for SARS-CoV-2 N and S genes, respectively. We observed similar partitioning behavior for SARS-CoV-2, RSV, RV, and MS2 RNA in our study; except for the partition of SARS-CoV-2 RNA at 22**°**C, which was higher (by approximately one order of magnitude) compared to other viruses and temperature conditions.

There are very limited studies in the literature on the effects of temperature on virus adsorption to particles. One study examined how temperature may influence the isotherm and kinetic adsorption process of viral genetic markers in wastewater solids: Yang et al.^11^ examined the kinetic adsorption of Phi6, MS2, T4, and Phix174 RNA in primary sludge at two temperature conditions (4**°**C and 25**°**C). They found that the rate of virus adsorption increased with increasing temperature (i.e., the time needed to reach equilibrium was reduced). Other studies have examined how temperature affects viral adsorption to clays. Syngouna et al. examined the isotherm and kinetic adsorption of infectious MS2 and ™X174 in clay particles at 4**°**C and 25**°**C, and found results similar to Yang et al.^31^ However, Bellou et al.^32^ measured the adsorption of MS2, ™X174, and hAdV RNA on clay particles at 4**°**C and 25**°**C and found that adsorption increased with decreasing temperature for hAdV, but decreased with decreasing temperature for MS2 and ™X174. In our isotherm (equilibrium) experiments, wastewater temperature did not seem to have an impact on the adsorption of viral genetic markers, except for the case of SARS-CoV-2. For SARS-CoV-2, a higher partition coefficient was observed at 22**°**C than at 4**°**C. The equivocal results described here suggest additional work is needed to better understand how temperature affects viral adsorption.

Limited previous research suggests that the presence of a lipid envelope outside the viral protein capsid of a virus may impact the solid-liquid partitioning of viruses and viral genetic markers in wastewater. For example, Ye et al.^12^ measured the adsorption of infectious MHV, ϕ6, MS2, and T3 in wastewater samples and found that enveloped viruses (MHV and ϕ6) were more strongly associated with solids than nonenveloped viruses (MS2 and T3). Similar results were reported in the study by Yang et al.,^11^ where they examined the solid-liquid partitioning behavior of Phi6, MS2, T4, and Phix174 RNA in primary sludge samples; researchers found that the majority of viral genetic markers were in the solid fraction of primary sludge. K_F_ ranged from 8,500–4,100,000 ml·g^-1^ for Phi6, MS2, T4, and Phix174 RNA. In our study, we observed that the partition and distribution behavior across enveloped (SARS-CoV-2 and RSV) and nonenveloped (RV and F+coliphage/MS2) viruses were the same. Our results were also similar across the partition and distribution experiments, which suggest that lab-grown viruses and endogenous viruses for SARS-CoV-2, RSV, RV, and F+coliphage/MS2 RNA may exhibit similar solid-liquid partitioning behavior in wastewater.

A few mechanistic-, statistical- and epidemiological-based models have been recently proposed to estimate the number of infected individuals within a sewershed. For example, Soller et al.^33^ developed a mechanistic model that estimates the fraction of a sewershed population actively infected with SARS-CoV-2. Wolfe et al.^34^ also developed a mass balance model that links SARS-CoV-2 RNA concentrations in solids to the number of individuals shedding SAR-CoV-2 RNA in stool within the sewershed. These models can potentially be applied to other viruses of interest however, a key parameter of these mechanistic models is the solid-liquid partitioning coefficient of viral genetic markers in wastewater solids. Limited information is available on viral-specific partitioning data, limiting the application of these models and the interpretation of wastewater surveillance data. Our results presented here fill a knowledge gap by providing information on virus partitioning that can be used in these modeling applications.

Understanding the fate and transport of viral genetic markers could inform wastewater sampling strategies and help optimize methods for processing wastewater and primary sludge samples. For example, a study by Kim et al^14^ showed that methods for processing influent and settled solids have comparable sensitivity. However, settled solids might be a more advantageous medium in sewersheds that have a low level of active infections because it requires less sample volume compared to influent methods. Our results also suggest that viral genetic markers might have similar partition/distribution coefficients across wastewater treatment plants. The results from our experiments might be applicable to other wastewater treatment plants that do not have local partition data available. Further research should be conducted to examine the solid-liquid partitioning of other viruses of interest for WBE efforts and evaluate how solid characteristics (e.g. particle size and biological/ chemical composition) might influence their partitioning behavior in wastewater.

### Environmental Implications

This study fills an important knowledge gap on the partitioning of respiratory viruses in wastewater and indicates that they partition preferentially to the solids.This information is useful in wastewater-based epidemiology applications (both sampling and modeling) but is also useful for informing wastewater treatment unit processes. These findings add to a growing body of evidence that the solids in wastewater (defined as material generally larger than 0.3 µm in hydrodynamic diameter) is enriched with infectious disease targets like viral RNA relative to the liquid phase on a mass-equivalent basis. (Note that intact bacteria and fungi, given their sizes, automatically fall into this size class.) There has been some confusion among researchers and practitioners in interpreting this statement. A volume of raw wastewater contains a small mass of solids (typically on the order of 10^2^ mg/L), so it can be true that most of the infectious disease target (mass or number) in that volume is in the liquid phase, and that the concentration of the infection disease target is enriched orders of magnitude in the solid relative to the liquid phase. The fact that infectious disease targets are enriched in the solid phase indicates that sample efforts that enrich for solids and choose the solids as a measurement matrix will improve the sensitivity of measurement approaches.

## Acknowledgments

This research was performed on the ancestral and unceded lands of the Muwekma Ohlone people. We pay our respects to them and their Elders, past and present, and are grateful for the opportunity to live and work here.

## References

(1) Boehm, A. B.; Hughes, B.; Doung, D.; Chan-Herur, V.; Buchman, A.; Wolfe, M. K.; White, B. J.Wastewater Surveillance of Human Influenza, Metapneumovirus, Parainfluenza, Respiratory Syncytial Virus (RSV), Rhinovirus, and Seasonal Coronaviruses during the COVID-19 Pandemic; preprint; Infectious Diseases (except HIV/AIDS), 2022. https://doi.org/10.1101/2022.09.22.22280218.

(2) Mercier, E.; D’Aoust, P. M.; Thakali, O.; Hegazy, N.; Jia, J.-J.; Zhang, Z.; Eid, W.; Plaza-Diaz, J.; Kabir, M. P.; Fang, W.; Cowan, A.; Stephenson, S. E.; Pisharody, L.; MacKenzie, E.; Graber, T. E.; Wan, S.; Delatolla, R. Municipal and Neighbourhood Level Wastewater Surveillance and Subtyping of an Influenza Virus Outbreak.Sci Rep 2022, 12 (1), 15777. https://doi.org/10.1038/s41598-022-20076-z.

(3) Ahmed, W.; Bivins, A.; Stephens, M.; Metcalfe, S.; Smith, W. J. M.; Sirikanchana, K.; Kitajima, M.; Simpson, S. L. Occurrence of Multiple Respiratory Viruses in Wastewater in Queensland, Australia: Potential for Community Disease Surveillance.Science of The Total Environment 2023, 864, 161023. https://doi.org/10.1016/j.scitotenv.2022.161023.

(4) Rector, A.; Bloemen, M.; Thijssen, M.; Pussig, B.; Beuselinck, K.; Van Ranst, M.; Wollants, E.Epidemiological Surveillance of Respiratory Pathogens in Wastewater in Belgium; preprint; Epidemiology, 2022. https://doi.org/10.1101/2022.10.24.22281437.

(5) Dumke, R.; Geissler, M.; Skupin, A.; Helm, B.; Mayer, R.; Schubert, S.; Oertel, R.; Renner, B.; Dalpke, A. H. Simultaneous Detection of SARS-CoV-2 and Influenza Virus in Wastewater of Two Cities in Southeastern Germany, January to May 2022.IJERPH 2022, 19 (20), 13374. https://doi.org/10.3390/ijerph192013374.

(6) Kirby, A. E.; Walters, M. S.; Jennings, W. C.; Fugitt, R.; LaCross, N.; Mattioli, M.; Marsh, Z. A.; Roberts, V. A.; Mercante, J. W.; Yoder, J.; Hill, V. R. Using Wastewater Surveillance Data to Support the COVID-19 Response — United States, 2020–2021.MMWR Morb. Mortal. Wkly. Rep. 2021, 70 (36), 1242–1244. https://doi.org/10.15585/mmwr.mm7036a2.

(7) Xagoraraki, I. Can We Predict Viral Outbreaks Using Wastewater Surveillance? J. Environ. Eng. 2020, 146 (11), 01820003. https://doi.org/10.1061/(ASCE)EE.1943-7870.0001831.

(8) Xagoraraki, I.; O’Brien, E. Wastewater-Based Epidemiology for Early Detection of Viral Outbreaks. In Women in Water Quality; O’Bannon, D. J., Ed.; Women in Engineering and Science; Springer International Publishing: Cham, 2020; pp 75–97. https://doi.org/10.1007/978-3-030-17819-2_5.

(9) Kitajima, M.; Ahmed, W.; Bibby, K.; Carducci, A.; Gerba, C. P.; Hamilton, K. A.; Haramoto, E.; Rose, J. B. SARS-CoV-2 in Wastewater: State of the Knowledge and Research Needs.Science of The Total Environment 2020, 739, 139076. https://doi.org/10.1016/j.scitotenv.2020.139076.

(10) Jin, Y.; Flury, M. Fate and Transport of Viruses in Porous Media. In Advances in Agronomy; Elsevier, 2002; Vol. 77, pp 39–102. https://doi.org/10.1016/S0065-2113(02)77013-2.

(11) Yang, W.; Cai, C.; Dai, X. Interactions between Virus Surrogates and Sewage Sludge Vary by Viral Analyte: Recovery, Persistence, and Sorption.Water Research 2022, 210, 117995. https://doi.org/10.1016/j.watres.2021.117995.

(12) Ye, Y.; Ellenberg, R. M.; Graham, K. E.; Wigginton, K. R. Survivability, Partitioning, and Recovery of Enveloped Viruses in Untreated Municipal Wastewater.Environ. Sci. Technol. 2016, 50 (10), 5077–5085. https://doi.org/10.1021/acs.est.6b00876.

(13) Armanious, A.; Aeppli, M.; Jacak, R.; Refardt, D.; Sigstam, T.; Kohn, T.; Sander, M. Viruses at Solid–Water Interfaces: A Systematic Assessment of Interactions Driving Adsorption.Environ. Sci. Technol. 2016, 50 (2), 732–743. https://doi.org/10.1021/acs.est.5b04644.

(14) Kim, S.; Kennedy, L.; Wolfe, M.; Criddle, C.; Duong, D.; Topol, A.; White, B. J.; Kantor, R.; Nelson, K.; Steele, J.; Langlois, K.; Griffith, J.; Zimmer-Faust, A.; McLellan, S.; Schussman, M.; Armmerman, M.; Wigginton, K.; Bakker, K.; Boehm, A.SARS-CoV-2 RNA Is Enriched by Orders of Magnitude in Solid Relative to Liquid Wastewater at Publicly Owned Treatment Works; preprint; Infectious Diseases (except HIV/AIDS), 2021. https://doi.org/10.1101/2021.11.10.21266138.

(15) Graham, K. E.; Loeb, S. K.; Wolfe, M. K.; Catoe, D.; Sinnott-Armstrong, N.; Kim, S.; Yamahara, K. M.; Sassoubre, L. M.; Mendoza Grijalva, L. M.; Roldan-Hernandez, L.; Langenfeld, K.; Wigginton, K. R.; Boehm, A. B. SARS-CoV-2 RNA in Wastewater Settled Solids Is Associated with COVID-19 Cases in a Large Urban Sewershed.Environ. Sci. Technol. 2021, 55 (1), 488–498. https://doi.org/10.1021/acs.est.0c06191.

(16) Li, B.; Di, D. Y. W.; Saingam, P.; Jeon, M. K.; Yan, T. Fine-Scale Temporal Dynamics of SARS-CoV-2 RNA Abundance in Wastewater during A COVID-19 Lockdown.Water Research 2021, 197, 117093. https://doi.org/10.1016/j.watres.2021.117093.

(17) Kitamura, K.; Sadamasu, K.; Muramatsu, M.; Yoshida, H. Efficient Detection of SARS-CoV-2 RNA in the Solid Fraction of Wastewater.Science of The Total Environment 2021, 763, 144587. https://doi.org/10.1016/j.scitotenv.2020.144587.

(18) Wolfe, M. K.; Duong, D.; Bakker, K. M.; Ammerman, M.; Mortenson, L.; Hughes, B.; Martin, E. T.; White, B. J.; Boehm, A. B.; Wigginton, K. R.Wastewater-Based Detection of an Influenza Outbreak; preprint; Public and Global Health, 2022. https://doi.org/10.1101/2022.02.15.22271027.

(19) Wolfe, M. K.; Yu, A. T.; Duong, D.; Rane, M. S.; Hughes, B.; Chan-Herur, V.; Donnelly, M.; Chai, S.; White, B. J.; Vugia, D. J.; Boehm, A. B. Use of Wastewater for Mpox Outbreak Surveillance in California.N Engl J Med 2023, 388 (6), 570–572. https://doi.org/10.1056/NEJMc2213882.

(20) Yin, Z.; Voice, T. C.; Tarabara, V. V.; Xagoraraki, I. Sorption of Human Adenovirus to Wastewater Solids.J. Environ. Eng. 2018, 144 (11), 06018008. https://doi.org/10.1061/(ASCE)EE.1943-7870.0001463.

(21) Hayes, E. K.; Sweeney, C. L.; Fuller, M.; Erjavec, G. B.; Stoddart, A. K.; Gagnon, G. A. Operational Constraints of Detecting SARS-CoV-2 on Passive Samplers Using Electronegative Filters: A Kinetic and Equilibrium Analysis.ACS EST Water 2022, pacsestwater.1c00441. https://doi.org/10.1021/acsestwater.1c00441.

(22) Shah, S.; Gwee, S. X. W.; Ng, J. Q. X.; Lau, N.; Koh, J.; Pang, J. Wastewater Surveillance to Infer COVID-19 Transmission: A Systematic Review.Science of The Total Environment 2022, 804, 150060. https://doi.org/10.1016/j.scitotenv.2021.150060.

(23) Borchardt, M. A.; Boehm, A. B.; Salit, M.; Spencer, S. K.; Wigginton, K. R.; Noble, R. T. The Environmental Microbiology Minimum Information (EMMI) Guidelines: QPCR and DPCR Quality and Reporting for Environmental Microbiology.Environ. Sci. Technol. 2021, acs.est.1c01767. https://doi.org/10.1021/acs.est.1c01767.

(24) Hejkal, T. W.; Wellings, F. M.; Lewis, A. L.; LaRock, P. A. Distribution of Viruses Associated with Particles in Waste Water.Appl Environ Microbiol 1981, 41 (3), 628–634. https://doi.org/10.1128/aem.41.3.628-634.1981.

(25) Huisman, J. S.; Scire, J.; Caduff, L.; Fernandez-Cassi, X.; Ganesanandamoorthy, P.; Kull, A.; Scheidegger, A.; Stachler, E.; Boehm, A. B.; Hughes, B.; Knudson, A.; Topol, A.; Wigginton, K. R.; Wolfe, M. K.; Kohn, T.; Ort, C.; Stadler, T.; Julian, T. R. Wastewater-Based Estimation of the Effective Reproductive Number of SARS-CoV-2.Environ Health Perspect 2022, 130 (5), 057011. https://doi.org/10.1289/EHP10050.

(26) Hughes, B.; Duong, D.; White, B. J.; Wigginton, K. R.; Chan, E. M. G.; Wolfe, M. K.; Boehm, A. B.Respiratory Syncytial Virus (RSV) RNA in Wastewater Settled Solids Reflects RSV Clinical Positivity Rates; preprint; Infectious Diseases (except HIV/AIDS), 2021. https://doi.org/10.1101/2021.12.01.21267014.

(27) Standard Operating Procedures for Interlaboratory and Methods Assessment of the SARS-CoV-2 Genetic Signal in Wastewater.

(28) Kalam, S.; Abu-Khamsin, S. A.; Kamal, M. S.; Patil, S. Surfactant Adsorption Isotherms: A Review.ACS Omega 2021, 6 (48), 32342–32348. https://doi.org/10.1021/acsomega.1c04661.

(29) Kantor, R. S.; Nelson, K. L.; Greenwald, H. D.; Kennedy, L. C. Challenges in Measuring the Recovery of SARS-CoV-2 from Wastewater.Environ. Sci. Technol. 2021, 55 (6), 3514– 3519. https://doi.org/10.1021/acs.est.0c08210.

(30) Kim, S.; Boehm, A. B. Wastewater Monitoring of SARS-CoV-2 RNA at K-12 Schools: Comparison to Pooled Clinical Testing Data.PeerJ 2023, 11, e15079. https://doi.org/10.7717/peerj.15079.

(31) Syngouna, V. I.; Chrysikopoulos, C. V. Interaction between Viruses and Clays in Static and Dynamic Batch Systems.Environ. Sci. Technol. 2010, 44 (12), 4539–4544. https://doi.org/10.1021/es100107a.

(32) Bellou, M. I.; Syngouna, V. I.; Tselepi, M. A.; Kokkinos, P. A.; Paparrodopoulos, S. C.; Vantarakis, A.; Chrysikopoulos, C. V. Interaction of Human Adenoviruses and Coliphages with Kaolinite and Bentonite.Science of The Total Environment 2015, 517, 86–95. https://doi.org/10.1016/j.scitotenv.2015.02.036.

(33) Soller, J.; Jennings, W.; Schoen, M.; Boehm, A.; Wigginton, K.; Gonzalez, R.; Graham, K. E.; McBride, G.; Kirby, A.; Mattioli, M. Modeling Infection from SARS-CoV-2 Wastewater Concentrations: Promise, Limitations, and Future Directions.Journal of Water and Health 2022, 20 (8), 1197–1211. https://doi.org/10.2166/wh.2022.094.

(34) Wolfe, M. K.; Archana, A.; Catoe, D.; Coffman, M. M.; Dorevich, S.; Graham, K. E.; Kim, S.; Grijalva, L. M.; Roldan-Hernandez, L.; Silverman, A. I.; Sinnott-Armstrong, N.; Vugia, D. J.; Yu, A. T.; Zambrana, W.; Wigginton, K. R.; Boehm, A. B. Scaling of SARS-CoV-2 RNA in Settled Solids from Multiple Wastewater Treatment Plants to Compare Incidence Rates of Laboratory-Confirmed COVID-19 in Their Sewersheds.Environ. Sci. Technol. Lett. 2021, 8 (5), 398–404. https://doi.org/10.1021/acs.estlett.1c00184.

(35) Montiel-Garcia, D.; Santoyo-Rivera, N.; Ho, P.; Carrillo-Tripp, M.; Iii, C. L. B.; Johnson, J. E.; Reddy, V. S. VIPERdb v3.0: A Structure-Based Data Analytics Platform for Viral Capsids.Nucleic Acids Research 2021, 49 (D1), D809–D816. https://doi.org/10.1093/nar/gkaa1096.

(36) Kim, S.; Kennedy, L. C.; Wolfe, M. K.; Criddle, C. S.; Duong, D. H.; Topol, A.; White, B. J.; Kantor, R. S.; Nelson, K. L.; Steele, J. A.; Langlois, K.; Griffith, J. F.; Zimmer-Faust, A. G.; McLellan, S. L.; Schussman, M. K.; Ammerman, M.; Wigginton, K. R.; Bakker, K. M.; Boehm, A. B. SARS-CoV-2 RNA Is Enriched by Orders of Magnitude in Primary Settled Solids Relative to Liquid Wastewater at Publicly Owned Treatment Works.Environ. Sci.: Water Res. Technol. 2022, 8 (4), 757–770. https://doi.org/10.1039/D1EW00826A.

(37) Wolfe, M. K.; Duong, D.; Bakker, K. M.; Ammerman, M.; Mortenson, L.; Hughes, B.; Arts, P.; Lauring, A. S.; Fitzsimmons, W. J.; Bendall, E.; Hwang, C. E.; Martin, E. T.; White, B. J.; Boehm, A. B.; Wigginton, K. R. Wastewater-Based Detection of Two Influenza Outbreaks.Environ. Sci. Technol. Lett. 2022, 9 (8), 687–692. https://doi.org/10.1021/acs.estlett.2c00350.

