## Supporting Information for "Adsorption of respiratory syncytial virus (RSV), rhinovirus, SARS-CoV-2, and F+ bacteriophage MS2 RNA onto wastewater solids from raw wastewater"

**Number of pages: 10**

**Number of Tables: 5**

**Number of Figures: 5**

#### Linear and Langmuir models

The Linear and Langmuir isotherm equations are defined as

Linear:: $$K_d = q_e/C_e \quad (1)$$

Langmuir: $$q_e = \frac{q_{max}K_L\,C_e}{1+\,K_LC_e} \quad (2)$$

where $q_e$ is the equilibrium concentration of viral genomes in solids (cp/g), $C_e$ is the equilibrium concentration of viral genomes in the liquid fraction of wastewater, $q_{max}$ is the maximum adsorption capacity (cp/g), $K_L$ is the Langmuir constant, and $K_d$ is the distribution coefficient.

#### Additional details related to EMMI[1] guidelines

The average number of droplets per two merged wells was 40,476 (standard deviation = 3,184). The machine vendor reports the droplet size as 0.00085 µL. Average number of copies per partition (λ) (standard deviation) for SARS-CoV-2 N gene was 0.027 (0.0008) and for BCoV was 0.238 (0.004). λ for other human viruses was similar to SARS-CoV-2.

**Table S1: Primers and probes for RT-ddPCR assays**

| Target | Sequence | Reference |
|---|---|---|
| SARS-CoV-2<br>(N gene) | Forward primer: 5′-CATTACGTTTGGTGGACCCT-3'<br>Reverse primer: 5′-CCTTGCCATGTTGAGTGAGA-3'<br>Probe: CGCGATCAAAACAACGTCGG (5' FAM/ZEN/3' IBFQ) | [2] |
| RSV<br>(N gene) | Forward primer: 5′- CTCCAGAATAYAGGCATGAYTCTCC-3'<br>Reverse primer: 5′- GCYCTYCTAATYACWGCTGTAAGAC-3'<br>Probe: TAACCAAATTAGCAGCAGGAGATAGATCAG (5' HEX/ZEN/3' IBFQ) | [3] |
| RV | Forward primer: 5′-GCCYGCGTGGCKGCC-3'<br>Reverse primer: 5′-GAAACACGGACACCCAAAG-3'<br>Probe: TCCTCCGGCCCCTGAATG (5' FAM/ZEN/3' IBFQ) | [3] |
| MS2 | Forward: 5'-GTCCATACCTTAGATGCGTTAGC-3'<br>Reverse: 5'-CCGTTAGCGAAGTTGCTTGG-3'<br>Probe: ACGTCGCCAGTTCCGCCATTGTCG (5' FAM/ZEN/3' IBFQ) | [4,5] |
| BCoV<br>(Extraction control) | Forward primer: 5’-CTGGAAGTTGGTGGAGTT-3’<br>Reverse primer: 5’-ATTATCGGCCTAACATACATC-3’<br>Probe: CCTTCATATCTATACACATCAAGTTGTT (5' FAM/ZEN/3' IBFQ) | [6] |

All probes contained fluorescent molecules and quenchers (5′ FAM and or HEX/ZEN/3′ IBFQ); FAM, 6-fluorescein amidite; HEX, hexachloro-fluorescein; ZEN, a proprietary internal quencher from Intrgrated DNA Technologies (Coralville, IA, USA); and IBFQ, Iowa Black FQ.

**Table S2: Thermal cycling conditions for SARS-CoV-2, RSV, RV, MS2, and BCoV**

| Cycling Step | Temperature °C | Time | Number of Cycles |
|---|---|---|---|
| Reverse transcription | 50 | 60 min | 1 |
| Enzyme activation | 95 | 10 min | 1 |
| Denaturation | 95 | 30 sec | 40 |
| Annealing/extension | SARS-CoV-2 and RSV: 59<br>RV: 61<br>MS2: 60<br>BCoV: 56 | 1 min* | 40 |
| Enzyme deactivation | 98 | 10 min | 1 |
| Hold | 4 | Infinite | 1 |

*Ramp rate set to 2°C/sec

**Table S3: Served population and annual average daily flow (MGD) of wastewater treatment plants**

| Plant | Location | Approximate number of people served in the sewershed† | Annual Average daily flow (MGD) | Min and Max TSS reported in 2022 (mg/L) | Min and Max pH reported in 2022 [-] |
|---|---|---|---|---|---|
| Oceanside Water Pollution Control Plant (OS) | San Francisco | 250,000 | 21.7 | 33–1,630 | * |
| Southeast Water Pollution Control Plant (SE) | San Francisco | 580,000 | 56.9 | 26–727 | 7.0–7.9 |
| Silicon Valley Clean Water Wastewater Treatment Plant (SV) | Redwood City | 199,000 | 12.4 | 160–400 | 7.0–7.6 |
| Sunnyvale Water Pollution Control Plant (SU | Sunnyvale | 161,021 | 12.6 | 164–376 | 6.4–7.2 |
| San Jose-Santa Clara Regional Wastewater Facility (SJ) | San Jose | 1,419,393 | 90.7 | 250–400 | 7.5–7.9 |
| South County Regional Wastewater Treatment Plant (GI) | Gilroy | 105,394 | 8.5 | 152–552 | 6.5–7.3 |

Notes:

† Based on the most recent National Pollution Discharge Elimination System (NPDES) permit and U.S. Census Bureau, 2020 American Community Survey block data

* Wastewater treatment plant does not measure pH in raw influent

**Table S4: Langmuir and Linear isotherm parameters for SARS-CoV-2, RSV-A, RV-B, and MS2 into wastewater solids**

| Virus | Temp. (°C) | Langmuir model | | Linear model |
|---|---|---|---|---|
| | | $K_L$ ($g \cdot ml^{-1}$) | $q_{max}$ ($cp \cdot g^{-1}$) | $K_d$ ($g \cdot ml^{-1}$) |
| SARS-CoV-2 | 4 | $4.5x10^{-4}$ | $2.6x10^{7}$ | 7,460 |
| | 22 | $1.0x10^{-3}$ | $9.5x10^{7}$ | 46,044 |
| RSV-A | 4 | $6.2x10^{-4}$ | $2.6x10^{8}$ | 136,809 |
| | 22 | $2.6x10^{-3}$ | $2.5x10^{8}$ | 425,372 |
| RV-B | 4 | $3.3x10^{-4}$ | $2.3x10^{7}$ | 3,396 |
| | 22 | $4.0x10^{-6}$ | $1.6x10^{7}$ | 33 |
| MS2 | 4 | $4.1x10^{-5}$ | $6.1x10^{7}$ | 1,260 |
| | 22 | $3.3x10^{-5}$ | $1.1x10^{7}$ | 148 |

**Table S5: Background concentration of endogenous of SARS-CoV-2, RSV, RV, and F+ coliphage/MS2 in wastewater influent samples stored at 4°C and 22°C**

| Virus | 4°C experiment | | 22°C experiment | |
|---|---|---|---|---|
| | Conc. in liquid fraction (cp/ml) | Conc. in solid fraction (cp/g) | Conc. in liquid fraction (cp/ml) | Conc. in solid fraction (cp/g) |
| **SARS-CoV-2** | 0.50 | 13,451 | 0.33 | 93,752 |
| **RSV** | 0.67 | 4,208 | 0.97 | 22,190 |
| **RV** | 0.88 | 90,729 | 0.87 | 22,580 |
| **F+ coliphage/ MS2** | 0.22 | 28,731 | 0.25 | 4,121 |

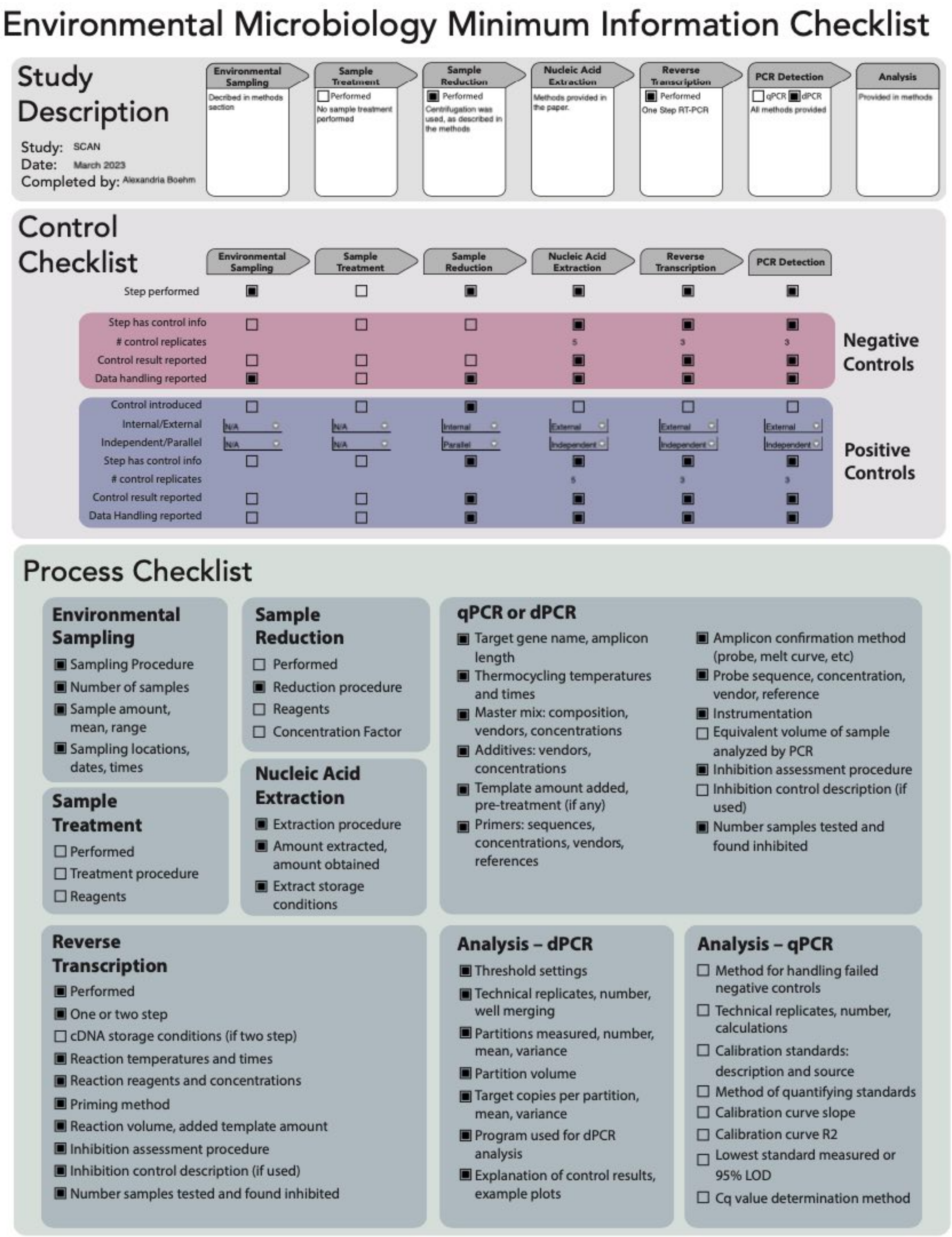


Figure S1. EMMI[1] checklist for reporting.

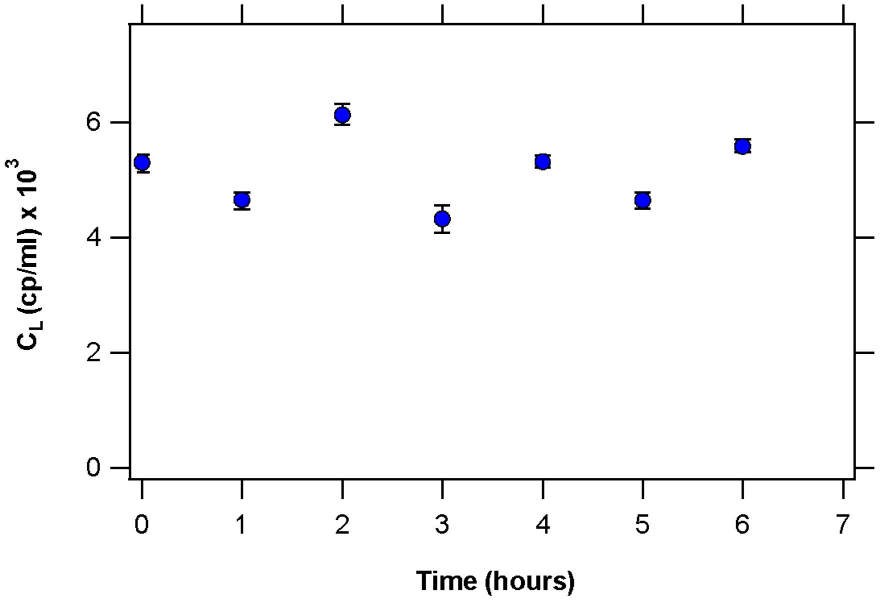


**Figure S2: MS2 RNA concentration in liquid fraction at t=0,1,2,3,4,5, and 6 hours. Error bars represent the 68% confidence interval from the ddPCR. Background concentration: ND**

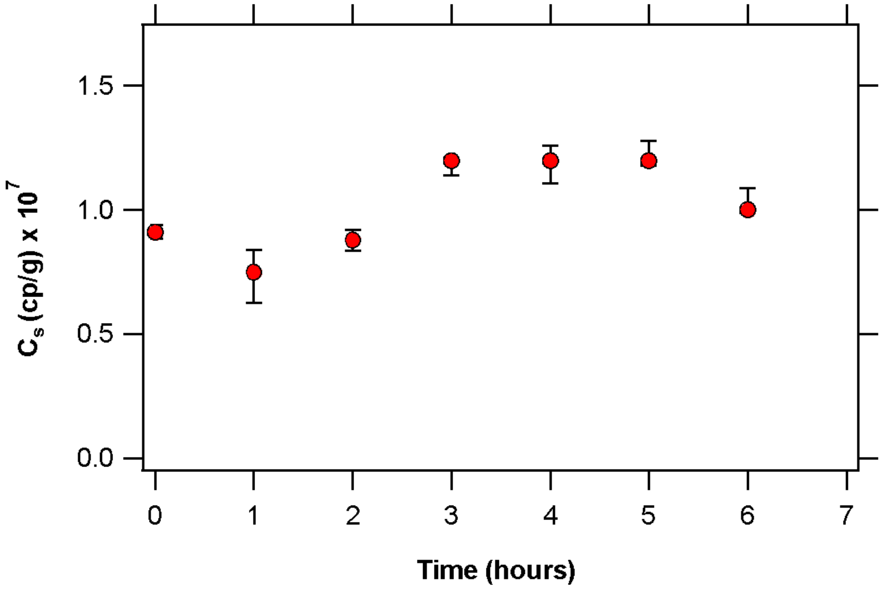


**Figure S3: MS2 RNA concentration in solid fraction at t=0,1,2,3,4,5, and 6 hours. Error bars represent the 68% confidence interval from the ddPCR. Background concentration ~ $3.6 \times 10^4$ gc/g dry.**

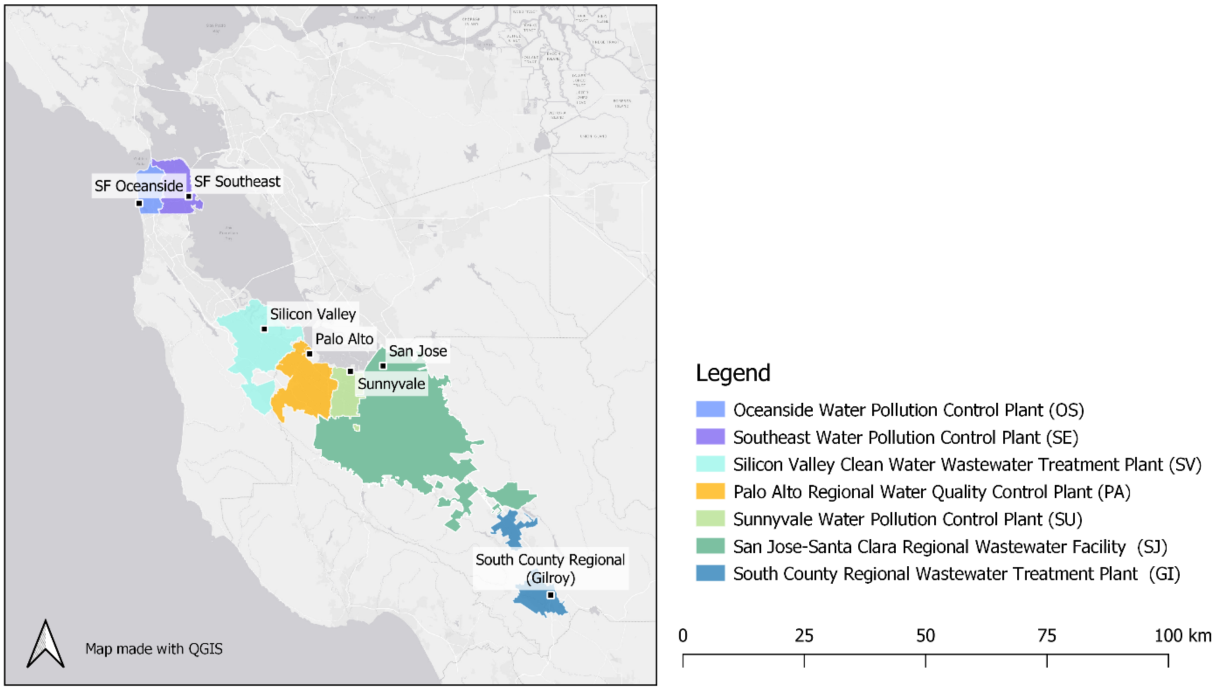


**Figure S4: Outline of wastewater treatment plant service area (sewersheds)**

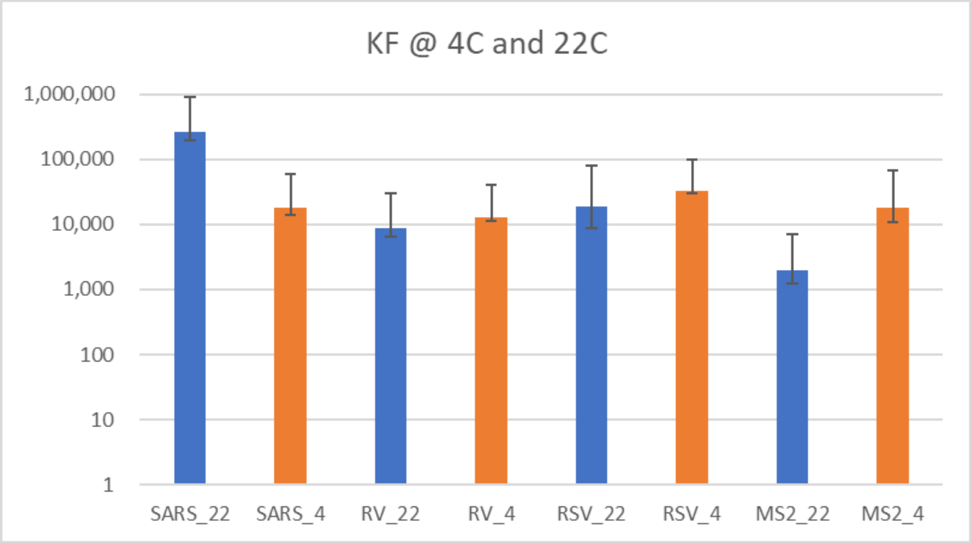


**Figure S5: Partition coefficient ($K_F$) for SARS-CoV-2, RV, RV, and MS2 in wastewater influent samples stored at 4°C and 22°C. Standard errors (SE) were obtained from the linear regression of the Freundlich model in R (summary function).**

### References

(1) Borchardt, M. A.; Boehm, A. B.; Salit, M.; Spencer, S. K.; Wigginton, K. R.; Noble, R. T. The Environmental Microbiology Minimum Information (EMMI) Guidelines: QPCR and DPCR Quality and Reporting for Environmental Microbiology. *Environ. Sci. Technol.* **2021**, acs.est.1c01767. https://doi.org/10.1021/acs.est.1c01767.

(2) Wolfe, M. K.; Topol, A.; Knudson, A.; Simpson, A.; White, B.; Vugia, D. J.; Yu, A. T.; Li, L.; Balliet, M.; Stoddard, P.; Han, G. S.; Wigginton, K. R.; Boehm, A. B. High-Frequency, High-Throughput Quantification of SARS-CoV-2 RNA in Wastewater Settled Solids at Eight Publicly Owned Treatment Works in Northern California Shows Strong Association with COVID-19 Incidence. *mSystems* **2021**, *6* (5), e00829-21. https://doi.org/10.1128/mSystems.00829-21.

(3) Boehm, A. B.; Hughes, B.; Duong, D.; Chan-Herur, V.; Buchman, A.; Wolfe, M. K.; White, B. J. Wastewater Concentrations of Human Influenza, Metapneumovirus, Parainfluenza, Respiratory Syncytial Virus, Rhinovirus, and Seasonal Coronavirus Nucleic-Acids during the COVID-19 Pandemic: A Surveillance Study. *The Lancet Microbe* **2023**, S266652472200386X. https://doi.org/10.1016/S2666-5247(22)00386-X.

(4) Gendron, L.; Verreault, D.; Veillette, M.; Moineau, S.; Duchaine, C. Evaluation of Filters for the Sampling and Quantification of RNA Phage Aerosols. *Aerosol Science and Technology* **2010**, *44* (10), 893–901. https://doi.org/10.1080/02786826.2010.501351.

(5) Turgeon, N.; Toulouse, M.-J.; Martel, B.; Moineau, S.; Duchaine, C. Comparison of Five Bacteriophages as Models for Viral Aerosol Studies. *Appl Environ Microbiol* **2014**, *80* (14), 4242–4250. https://doi.org/10.1128/AEM.00767-14.

(6) Decaro, N.; Elia, G.; Campolo, M.; Desario, C.; Mari, V.; Radogna, A.; Colaianni, M. L.; Cirone, F.; Tempesta, M.; Buonavoglia, C. Detection of Bovine Coronavirus Using a TaqMan-Based Real-Time RT-PCR Assay. *Journal of Virological Methods* **2008**, *151* (2), 167–171. https://doi.org/10.1016/j.jviromet.2008.05.016.
